# An implant for long-term cervical vagus nerve stimulation in mice

**DOI:** 10.1101/2020.06.20.160473

**Authors:** Ibrahim T. Mughrabi, Jordan Hickman, Naveen Jayaprakash, Eleni S. Papadoyannis, Adam Abbas, Yao-Chuan Chang, Sunhee Lee, Timir Datta-Chaudhuri, Eric H. Chang, Theodoros P. Zanos, Robert C. Froemke, Cristin Welle, Yousef Al-Abed, Stavros Zanos

## Abstract

Vagus nerve stimulation (VNS) is a neuromodulation therapy with the potential to treat a wide range of chronic conditions in which inflammation is implicated, including type 2 diabetes, obesity, atherosclerosis and heart failure. Many of these diseases have well-established mouse models but due to the significant surgical and engineering challenges that accompany a reliable interface for long-term VNS in mice, the therapeutic implications of this bioelectronic approach remain unexplored. Here, we describe a long-term VNS implant in mice, developed at 3 research laboratories and validated for between-lab reproducibility. Implant functionality was evaluated over 3-8 weeks in 81 anesthetized or conscious mice by determining the stimulus intensity required to elicit a change in heart rate (heart rate threshold, HRT). HRT was also used as a method to standardize stimulation dosing across animals. Overall, 60-90% of implants produced stimulus-evoked physiological responses for at least 4 weeks, with HRT values stabilizing after the second week of implantation. Furthermore, stimulation delivered through 6-week-old implants decreased TNF levels in a subset of mice with acute inflammation caused by endotoxemia. Histological examination of 4- to 6-week-old implants revealed fibrotic encapsulation and no gross fiber loss. This implantation and dosing approach provide a tool to systematically investigate the therapeutic potential of long-term VNS in chronic diseases modeled in the mouse, the most widely used vertebrate species in biomedical research.

## Introduction

The vagus is a cranial nerve tasked with maintaining internal organ homeostasis by mediating several autonomic reflexes between the brain and periphery [1]. It also plays a central role in controlling inflammation via a recently mapped neuroimmune mechanism, termed the inflammatory reflex [2-5]. Work by Tracey et al. has demonstrated that vagus nerve signals decrease inflammatory cytokine release by macrophages in the spleen [6-8]. This finding stimulated a large body of work looking to expand the clinical utility of vagus nerve stimulation (VNS), an intervention already approved for epilepsy [9] and depression [10], to diseases with an inflammatory component. Currently, long-term VNS is being explored in the treatment of brain disorders, such as tinnitus [11, 12], stroke [13, 14], and Alzheimer’s disease [15, 16], as well as peripheral organ and systemic diseases, including heart failure [17], cardiac arrhythmias [18-21], pulmonary hypertension [22, 23], rheumatoid arthritis [24], Crohn’s disease [25], and lupus [26]. Additional possible indications for VNS include common disorders in which inflammation is implicated, such as type 2 diabetes, obesity, and atherosclerosis [27-30].

The preclinical study of VNS in models of chronic diseases requires a long-term VN implant and has been mostly limited to neurological and cardiovascular diseases modeled in rats and large animals [19, 21, 23, 31-38]. Although the mouse is considered the species of choice in the study of disease mechanisms and the standard for preclinical therapeutic screening [39, 40], translational VNS research in mice has been limited to acute delivery of stimulation [41-49]. This is mainly due to the significant surgical and technical challenges for a mechanically-[50-52] and electrochemically-[53, 54] stable long-term interface with the microscopic anatomy of the mouse vagus nerve. As a consequence, the therapeutic role of VNS in many models of chronic diseases is largely unexplored, as are the long-term effects of VNS. A functional and reliable long-term VNS implant in the mouse will broaden the translational potential of VNS and provide an experimental tool to probe the chronic effects of vagal neuromodulation.

Here we describe a surgical technique to permanently implant a micro-cuff electrode onto the mouse cervical vagus for long-term neurostimulation. The technique was developed and refined through a collaboration between 3 research labs (Feinstein Institutes, University of Colorado, and New York University) and validated in cohorts tested in 2 of those labs. We also provide a standardized method to test implant functionality in anesthetized or conscious animals and to deliver a consistent “stimulation dose” within and across animals over several weeks after implantation. In 4 cohorts of animals (81 mice in total), we document predictable and consistent longitudinal implant performance, with expected stimulus-elicited physiological responses for more than 4 weeks. We also demonstrate that VNS delivered through 6-week old implants was successful at reducing serum TNF levels in a subset of mice with acute endotoxemia. Finally, we find that implantation of these cuffs induced fibrotic encapsulation, but nerve fibers were largely preserved. This method will allow screening of VNS therapies in several models of disease in which inflammation is implicated and facilitate mechanistic studies of long-term vagal neuromodulation. Further, it will provide a tool to better understand the role of autonomic afferent signaling in memory and behavior.

## Methods

### Electrode preparation

Bipolar platinum-iridium micro-cuff electrodes with an internal diameter of 150 μm (MicroLeads Neuro, Somerville, MA) or 100 μm (CorTec, Germany) were soldered to gold sockets after cutting the lead wires to a length of 2.5-3.0 cm (Fig 1A). Electrical impedance was measured in saline for each electrode at 1 kHz using MicroProbes Impedance Tester (MicroProbes, Gaithersburg, MD). For sterilization, the soldered electrodes were submerged in 0.55% ortho-phthaladehyde solution (Cidex OPA, Advanced Sterilization Products, Irvine, CA) or 70% ethanol for 15 mins, rinsed 4 times with sterile saline, and if Cidex used sonicated in saline for 5 minutes as a final rinse.

**Fig 1.**
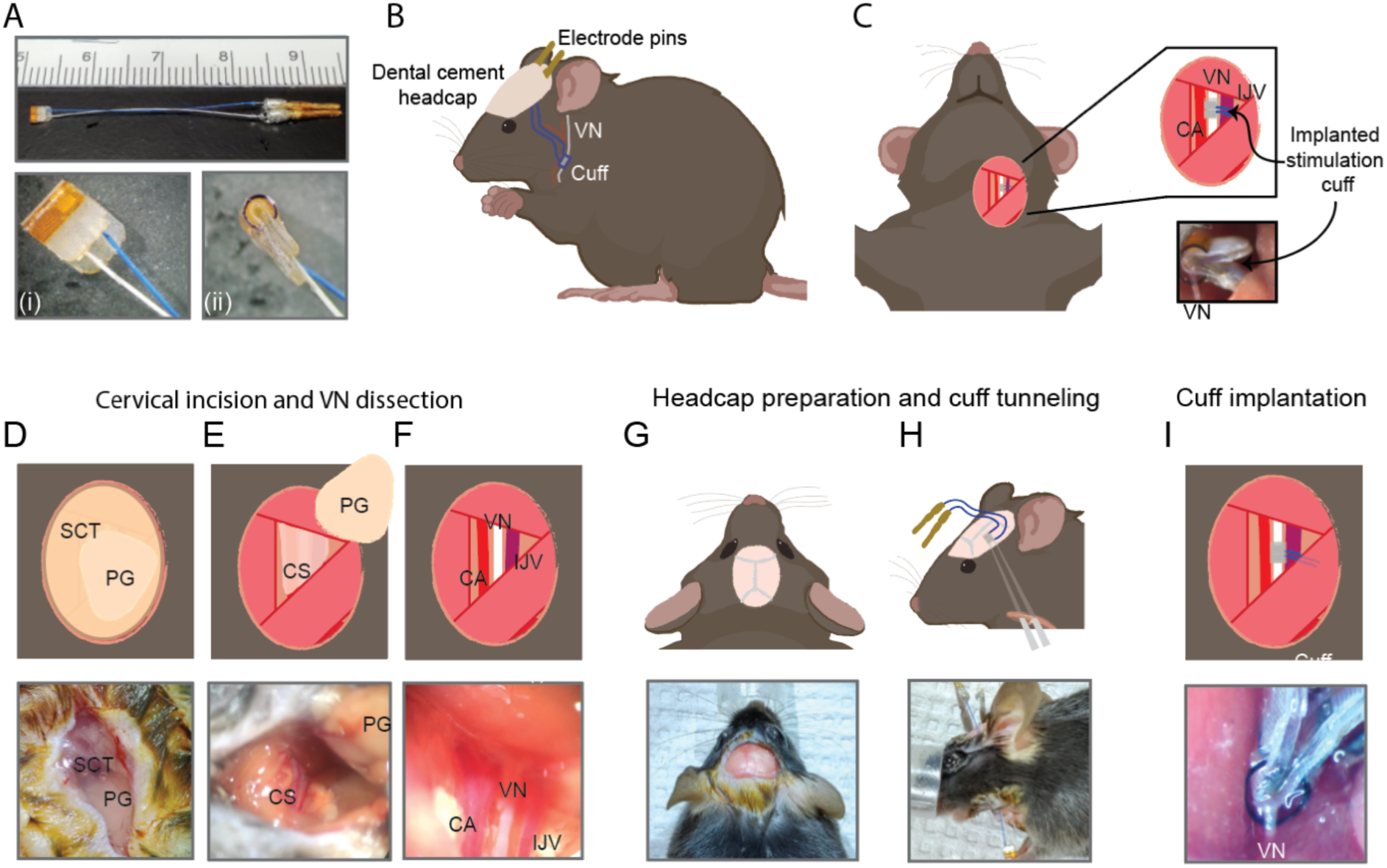
Surgical procedure for long-term implantation. (A) Lead wires of cuff electrodes were cut to a length of 2.5-3.0 cm and soldered to gold pins. Inset: en face (i) and side (ii) view of 150 μm Microleads cuff electrode. (B, C) Overview of implant, headcap with pins and location of the vagus nerve (VN) cuff. (D) A 1-cm ventral incision is made about 0.5 cm lateral to the sternal notch, exposing subcutaneous tissue (SCT) and the parotid gland (PG). (E) SCT is bluntly dissected freeing the PG which is then retracted from view exposing the carotid sheath (CS). (F) The vagus nerve (VN) is bluntly dissected away from the carotid artery (CA) and the internal jugular vein (IJV). (G) The scalp is incised to expose lambda and bregma. (H) A subcutaneous tunnel is created from skull base to cervical incision site, either between the eye and ear (depicted) or directly caudal to the ear. (I) The cuff is tunneled under the sternomastoid muscle and implanted on the vagus nerve. Pins are finally secured to the skull with dental cement. Created with BioRender.com.

### Implantation procedure

Male C57BL/6 mice were purchased from Charles River Laboratories (Wilmington, MA) at the age of 8-12 weeks. Animals were housed under 12-hour light/dark cycle with ad libitum access to food and water. All animal experiments complied with relevant ethical guidelines and were approved by the Institutional Animal Care and Use Committee (IACUC) of the Feinstein Institutes for Medical Research and University of Colorado Anschutz Medical Campus. The surgery features an implantation of a cuff electrode on the cervical vagus nerve with lead wires that are tunneled to a headcap secured with dental cement to the skull (Figs 1B and 1C).

Mice were placed on a heated surgical platform equipped with a dissecting microscope in the supine position under isoflurane anesthesia at 4% induction and 1.5% maintenance. Hair over the neck area was removed using a depilatory cream (Nair, Church & Dwight, Ewing, NJ) or with an electric shaver and skin disinfected using alternating swabs of betadine and 70% ethanol. A 1-cm vertical incision was made starting at the level of the sternal notch about 0.5 cm left of midline (Fig 1D). The left lobe of the parotid (salivary) gland (PG) was bluntly dissected away from subcutaneous tissue (SCT) (Figs 1D and 1E). The muscles forming the anterior triangle were exposed providing a window to view the carotid sheath (CS) housing the carotid artery (CA), vagus nerve (VN), and internal jugular vein (IJV). Using blunt dissection, the carotid sheath was isolated and the VN carefully dissected away from the fibrous connective tissue (Fig 1F). The incision site was temporarily closed, and the mouse was turned to the prone position to access the skull. The head was shaved and disinfected with alternating swabs of betadine and 70% ethanol. After application of a local analgesic, a fold of skin over the center of the skull was removed exposing bregma and lambda. The exposed skull was treated with 3 alternating rinses of hydrogen peroxide and saline, while gently scoring the skull surface with a sterile scalpel in between rinses (Fig 1G). After drying the skull with compressed air, acrylic dental cement (Metabond, Parkell, Edgewood, NY) was applied to the right half of the exposed area and allowed to set for 5 minutes. Using blunt dissection at the left edge of the scalp incision, a subcutaneous tunnel was created between the animal’s eye and ear down to the ventral neck incision location (Fig 1H). Alternatively, the subcutaneous tunnel can be performed from caudal to the dorsal ear at the base of the skull to the ventral incision. The pre-formed subcutaneous tunnel was accessed using blunt dissection at the lower edge of the neck incision, and a pair of fine straight forceps were used to pull the electrode cuff subcutaneously into the neck area (Fig 1H). The cuff was further tunneled under the sternomastoid muscle and placed on the VN situated close to the nerve’s original anatomic position with care taken to avoid excessive manipulation (Fig 1I). To confirm successful electrode placement, a brief stimulus was delivered through the externalized electrode leads to measure heart rate response. The neck incision was then closed with 6-0 nylon suture and the animal turned to the prone position. The externalized connectors were held at an angle against the exposed part of the skull and acrylic dental cement (Metabond) applied covering the leads and part of the connectors and allowed to set for 5 minutes (Fig 1B). To seal the headcap to the skin, liquid surgical adhesive (Vetbond, 3M, Saint Paul, MN) was applied to the skin-cement junction. The mice were moved to clean, warmed cages and monitored until conscious and mobile. The surgical procedures were carried out under strict aseptic conditions, and animals were supplemented with warm saline intra- and post-operatively.

In some experiments, mice were instrumented with implanted ECG electrodes to measure HRT in awake behaving animals. Following the surgical approach described above, 3 platinum wires were tunneled subcutaneously along the cuff leads from the skull to the ventral neck. The left ECG lead was tunneled subcutaneously through a 1-cm incision at the left costal margin and the exposed part fixed to the underlying muscle with 6-0 nylon suture. The right ECG lead was tunneled subcutaneously from the neck incision and sutured to the pectoralis muscle. The ground ECG lead was imbedded in the neck between the right lobe of the salivary gland and the skin. The ECG and cuff leads were connected to a multi-channel nano-connector (Omnetics Connector Corporation, Minneapolis, MN) and cemented to the skull as described before.

### Nerve stimulation and physiological monitoring

Experiments were carried out in 4 separate cohorts of mice at two independent laboratories: cohorts 1-3 at Feinstein Institutes, and cohort 4 at University of Colorado. In cohorts 1-3, electrode functionality was evaluated based on the ability to induce a decrease in heart rate (HR) during stimulation in anesthetized animals. Heart rate threshold (HRT) was defined as the minimum current intensity required to elicit a 5-15% drop in heart rate using a stimulus train of 300 bi-phasic, charge-balanced, square pulses at a pulsing frequency of 30 Hz with 0.1, 0.5, or 1 ms pulse widths (PW). In most cases, HRT was initially determined with 100 μs PWs, which was changed to 500 μs and finally to 1000 μs whenever HRT exceeded 2 mA; in 4 mice, HRT was determined at all 3 PWs over several sessions. In cohort 2, mice were tested on 3-7 days during the first week post-implantation then once or twice weekly thereafter, whereas cohorts 1 and 3 were tested less frequently or regularly. During testing sessions, anesthetized mice were instrumented with ECG electrodes and a nasal temperature sensor to measure ECG and nasal air flow and calculate heart and breathing rates. The physiological signals were amplified using a biological amplifier (Bio-amp Octal, ADInstruments) for ECG and Temperature Pod (ADInstruments) for nasal temperature and digitized using PoweLab 16/35 (ADInstruments). The digital signals were then streamed to a PC running LabChart v8 (ADInstruments). VNS was delivered by a rack-mounted stimulus generator (STG4008, Multichannel Systems, Reutlingen, BW Germany). In cohort 4, performed at University of Colorado Anschutz Medical Campus, stimulation response was defined as a reduction in HR measured with an infrared paw sensor (Mouse Stat Jr, Kent Scientific) or respiratory rate (measured visually) in response to a stimulus train of 0.2-1 mA intensity, 100us PW, and 30Hz frequency. Stimulation failure occurred when there was no response in either heart rate reduction or breathing rate alterations. Further failures included headcap failure. This cohort was designed for behavior and was tested regularly within the first 14 days. Thereafter, a random subset of the mice had additional stimulation testing on a per needed basis for further experiments.

In awake experiments, animals implanted with ECG leads were gently restrained and connected to a commutator (P1 Technologies, Roanoke, VA) that interfaced with the stimulus generator and the bio-amplifier; HRT was determined as described above. Intensity at maximum charge injection capacity (CIC) was calculated using the average reported value of CIC for platinum iridium (50 - 150 μC/cm^2^) [55, 56] applied to the implanted electrode surface area (0.00474 cm^2^) for PWs of 500 μs (cohort 2 and 3) and 600 μs (cohort 1).

### LPS endotoxemia challenge

Eight-week old mice were implanted with a left VN cuff as described before. HRT was determined once weekly for 4 weeks, then animals were left unstimulated for 2 weeks to avoid any long-lived VNS effects. On week 6 post-implantation, VNS was delivered to animals under anesthesia at HRT intensity using 250 ms PW and 10 Hz frequency for 5 minutes. Lipopolysaccharide (LPS, Sigma-Aldrich, St. Louis, MO) was administered to mice (0.1 mg/kg, i.p.) 3 hours after stimulation. Blood was collected by cardiac puncture 90 minutes post-LPS injection and left to clot for 1 hour at room temperature. The blood samples were then centrifuged at 2000 xg for 10 minutes and serum collected for TNF determination by ELISA (Invitrogen, Carlsbad, CA) following the manufacturer’s instructions.

### Histology and immunohistochemistry

Mice with chronic implants of at least 4 weeks were euthanized and segments of the neck were excised and fixed in 10% buffered formalin for at least 2 weeks. Fixed segments were then prepared for frozen sectioning. Serial cross-sections of the tissue specimens were obtained at 50 µm thickness using a cryostat. Standard immunohistochemical protocols were followed to stain the mounted sections for neurofilament [57]. Briefly, sections were rinsed with 1x Tris-buffered saline (TBS) then blocked for 1 hour using 1% normal goat serum and Triton X-100 (Sigma Aldrich) in TBS. Sections were then incubated with primary anti-bodies against neurofilament (1:500, ab8135, Abcam) overnight at 4°C. The following day, sections were rinsed and incubated with goat anti-rabbit Alexa 488 secondary antibody (1:500, Fisher Scientific) for two hours at room temperature. Following incubation, stained slides were rinsed 3 times with TBS buffer then mounted with Fluoromount-G (Thermo Fisher Scientific). Images of the vagus nerve were obtained with 100X magnification using a Keyence BZ-X810 fluorescence microscope (Keyence, Japan). The left vagus nerve was identified either within the tissues covering the upper margin of the cuff or in the most anterior part of the neck adjacent to the cuff. The right vagus nerve was identified in its anatomic position at the level of the esophagus. Fibers larger than 1.5 μm were measured and counted using ImageJ.

## Results

### Cervical vagus nerve stimulation through the long-term implant produces changes in heart rate and breathing rate

The cervical vagus nerve (VN) comprises parasympathetic motor and visceral sensory fibers that regulate many physiological functions, including heart rate (HR) and breathing [1, 58, 59]. To determine whether the implanted electrodes were functional, we stimulated the VN with increasing current intensity while measuring stimulus-elicited changes in HR and breathing rate (BR) in animals under isoflurane anesthesia. Cervical VNS produced decreases in HR as well as changes in BR (Fig 2A). The magnitude of HR reduction was dependent on current intensity (Fig 2B), whereas BR showed more variable responses, including tachypnea, bradypnea, and apnea (Figs 2A and 3A). In general, HR response was more reliable and occurred at lower intensities than BR response (data not shown). In awake behaving mice (n=2), VNS produced comparable dose-dependent HR responses (S1 Movie). Animals receiving awake VNS did not show any signs of distress or visible changes in BR.

**Fig 2.**
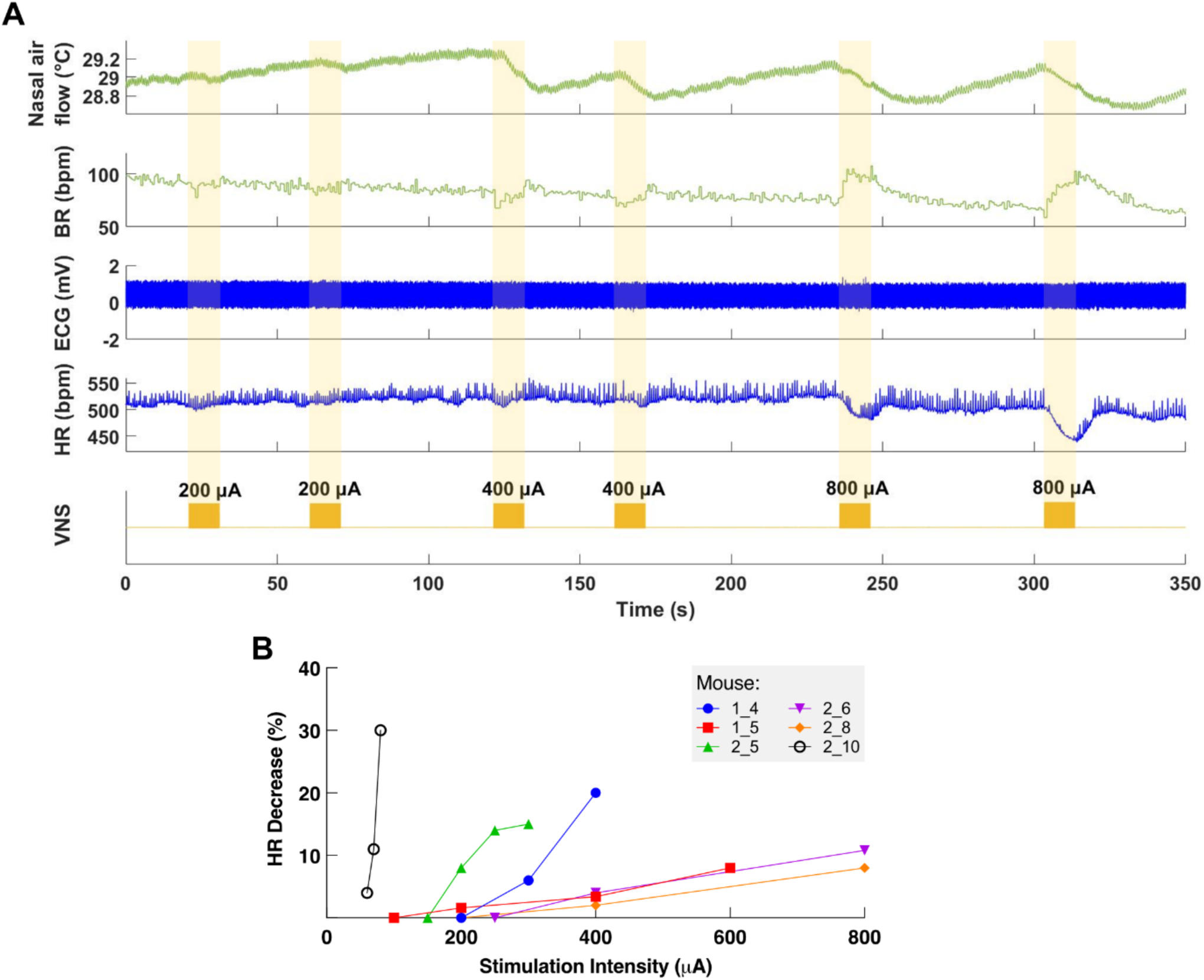
Dose-dependent physiological responses to cervical VNS. (A) Representative traces from a chronically implanted mouse showing nasal air flow (top panel) and extracted breathing rate (BR, second panel), and ECG (third panel) and extracted heart rate (HR, fourth panel). Trains of VNS of increasing intensity from 200 to 800 μA (fifth panel, yellow traces), caused BR and HR responses with increasing magnitudes. (B) Percentage of HR decrease as a function of VNS intensity in 6 chronically implanted mice.

### Longitudinal changes in heart rate threshold and electrode impedance

In order to assess the longitudinal functionality of each implant, we determined heart rate threshold (HRT) over time, defined as the minimum current intensity of a stimulus train (300 pulses at 30 Hz) required to elicit an approximately 5-15% decrease in HR. In cohorts 1-3 (completed in April, June, and October 2019 at the Feinstein Institutes for Medical Research) HRT and impedance values were determined for 3-8 weeks post-implantation. VNS elicited drops in HR and changes in breathing for up to 8 weeks post-implantation (Fig 3A). In mice of cohort 2, where HRT was measured more consistently, initial HRT values determined with 100 μs long pulses were variable among animals (range = 30 μA - 300 μA, mean = 102, SD = 88) and increased over the first week post-implantation (Pearson r = 0.52, *p* < 0.01), whereas HRT values determined with 500 μs long pulses did not change significantly with time (r = 0.04, *p* NS) (Fig 3B). HRT values at 100 μs PW were 54% greater on average than those at 500 μs PW, and that relationship was maintained over time (Fig 3C). Bipolar electrical impedance in cohort 2 decreased during the first 2 weeks (r = −0.49, *p* < 0.01) and then stabilized (r = 0.26, *p* = 0.08) (Fig 3D), while it remained relatively stable over time in cohorts 1 (r = 0.12, *p* = 0.29) and 3 (r = 0.39, *p* = 0.12) (S2 Fig). Pre-implantation impedance values did not correlate with initial HRT (r = 0.50 for cohort 1, r = 0.30 for cohort 2, *p* NS in both cases). Interestingly, there was no correlation between the changes in electrical impedance values and HRT (r = 0.05, *p* = 0.66); in some mice, non-functional cuffs continued to record relatively low impedance values despite their inability to induce a physiological response (Fig 3E). In cohorts 1-3, implant failure occurred more frequently in earlier compared to later cohorts. The percentage of mice with functional implants at 4 weeks post-implantation increased from 40% in the cohort 1 to 90% in cohort 3 (Table 1). Cohort 4 was specifically designed for behavioral experiments and performed at the University of Colorado Anschutz Medical Campus. All implanted mice were checked for stimulation efficacy and survival of the surgery within days 1-5 and had a success rate of 96% (50/52 mice). Mice that had successful stimulation following days 1-5 were then tested prior to behavioral experiments in days 6-14 and demonstrated a success rate of 76% (38/52) (Table 1). After this point, mice had stimulation checks performed on a per needed basis for further behavioral testing. Of these, 17/18 and 11/13 responded to stimulation in the 15-30 and 30+ days’ time period, respectively (Table 1). Implanted mice that were responsive to stimulation prior to behavioral testing remained responsive at a high rate, pointing to the long-term viability of the cuff implantation In all 4 cohorts, electrode failures generally occurred during the second week post-implantation and implant functionality stabilized thereafter (Table 1).

**Table 1.** Number of functional implants in each cohort across time.

| Days post-implantation: | Functional cuffs |  |  |  |
| --- | --- | --- | --- | --- |
|  | 1-5 | 6-14 | 15-29 | 30+ |
| **Cohort 1 (n=10)** | 8/10 (80%) | 6/10 (60%) | 4/10 (40%) | 4/10 (40%) |
| **Cohort 2 (n=10)** | 10/10 (100%) | 7/10 (70%) | 7/10 (70%) | 6/10 (60%) |
| **Cohort 3 (n=9)** | 9/9 (100%) | 8/9 (90%) | 8/9 (90%) | -- |
| **Cohort 4 (n=52)** | 50/52 (96%) | 38/52 (76%) | 17/18 (*) | 11/13 (*) |

\* Group is a random subset of the (6-14 days) functional implants (n=38).
Implants were tested in 4 cohorts. In cohorts 1-3, implant functionality was determined based on heart rate threshold, and implant failure was defined as the absence of a physiological response upon stimulation with 3 mA or higher on 3 consecutive testing sessions. In cohort 4, functionality was determined based on a reduction in heart rate or breathing rate and failure was defined as absence of response with 1 mA on 1 occasion.

**Fig 3.**
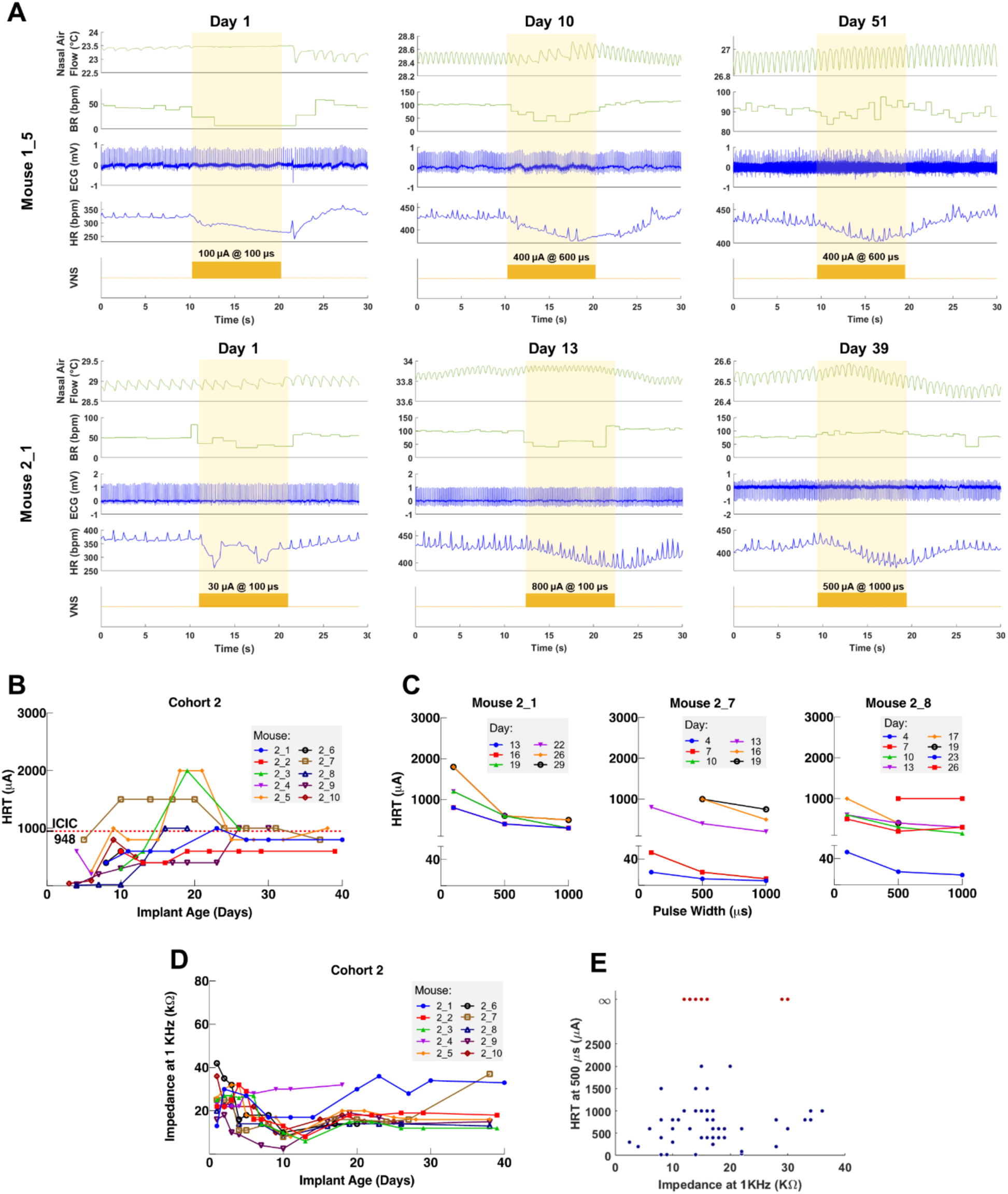
Longitudinal changes in heart rate threshold and electrode impedance. (A) Examples of threshold and supra-threshold responses at different implant ages in 2 mice. Heart rate threshold (HRT) was defined as the stimulation intensity required to produce a 5-15% decrease in HR. Mice elicited HR responses up to 2 months post-implantation. (B) HRT values vs. implant age in 10 mice from cohort 2. The horizontal red dotted line indicates intensity at maximum charge injection capacity (ICIC) calculated for the micro-cuff electrodes. (C) HRT values determined with VNS trains of 0.1, 0.5 and 1 ms-wide pulses at different implant ages, in 3 mice from cohort 2. (D) Electrical impedance values at 1KHz vs. implant age in mice from cohort 2. (E) HRT values plotted against electrode impedances in cohort 2 animals, from individual measurements performed during a 40-day period post-implant. Infinity HRT values (red-colored data points) indicate implants that did not produce a heart rate response up to 5mA. Pearson correlation was 0.05 (p NS).

### Implant is associated with fibrosis and preserved nerve fibers in the cuffed nerve

Long-term efficacy of peripheral nerve implants could deteriorate due to nerve damage and/or fibrous tissue encapsulation [60, 61]. To determine the impact of these processes in our long-term implants, we collected both vagus nerves, cuffed (left) and non-cuffed (right), from animals along with surrounding tissue at 4-6 weeks post-implantation for gross and histological analysis. The implant site exhibited moderate fibrosis encompassing the cuff surfaces in animals with both functional and non-functional implants (Fig 4A). Histological analysis revealed largely preserved nerve fibers in the cuffed compared with the non-cuffed nerve in the same animal with no obvious axon pathology or fragmentation (Fig 4B, S3 Fig).

**Figure 4.**
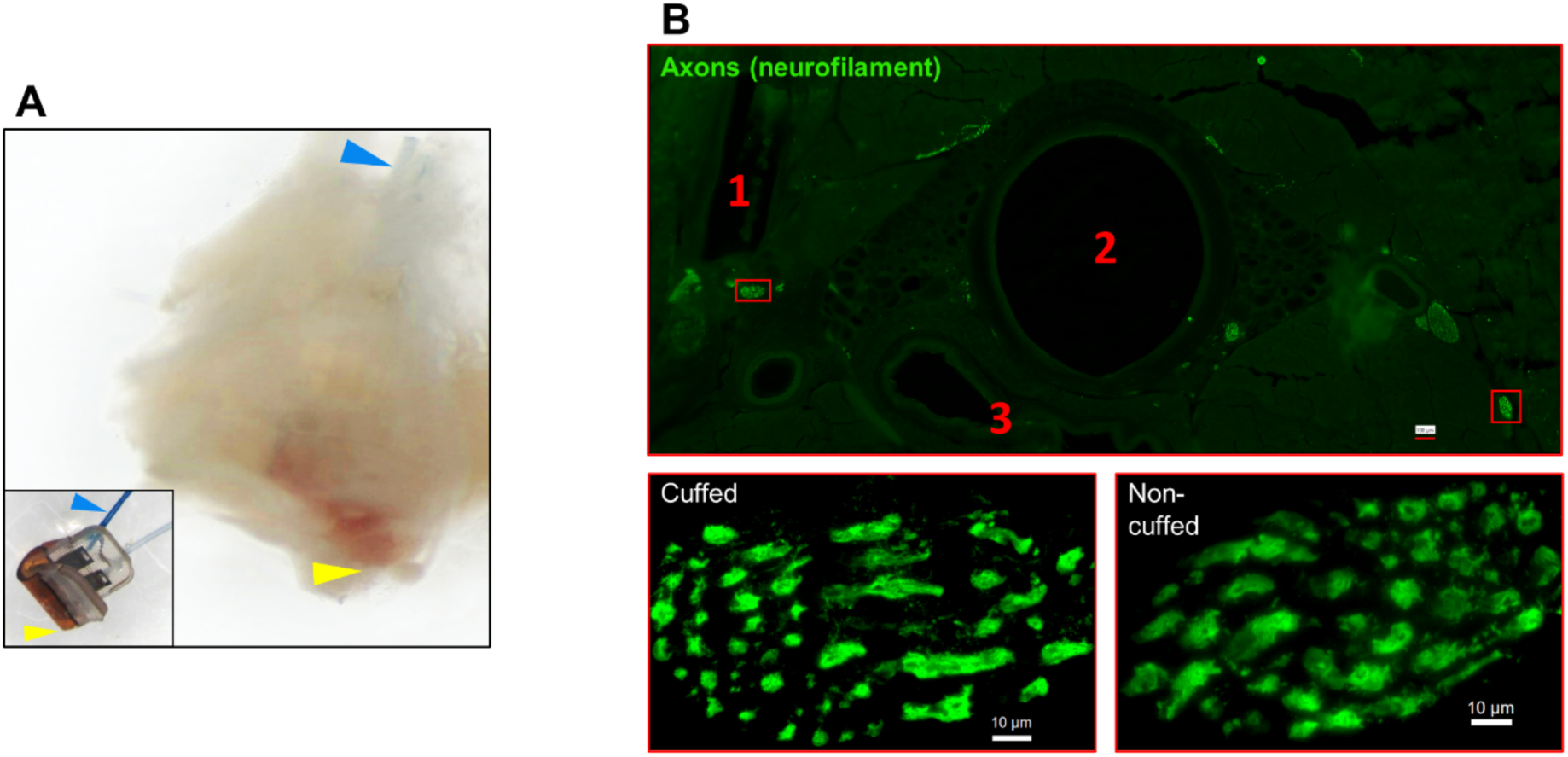
Gross anatomy and histology of cuffed nerve and surrounding tissues. (A) Left VN with a micro-cuff electrode dissected from fixed tissue, obtained from a mouse with a 6-week old implant, showing fibrotic encapsulation. Inset: micro-cuff electrode in a similar orientation, with highlighted lead (blue arrowhead) and cuff tip (yellow arrowhead). (B) Cross-section of the neck immediately rostral to the cuff (upper panel) from a mouse with a 6-week old implant stained for axons (neurofilament, green) and showing landmarks: (1) rostral cuff margin; (2) trachea; (3) esophagus. The cuffed left VN (magnified, lower left panel) is located adjacent to the cuff and shows 47 large fibers. The non-cuffed right VN (magnified, lower right panel) lies at the level of the esophagus and shows 43 large fibers.

### Functional 6-week old implants inhibit TNF release in some endotoxemic mice

Acute VNS decreases serum TNF levels in acute inflammation models by modulating the immune response [6]. To test whether our long-term implant can produce a similar effect, we used it to deliver one-time VNS in an LPS endotoxemia model (Fig 5A). Seven mice with 6-week old implants received VNS (intensity at HRT, PW 250 μs, frequency 10 Hz) or sham stimulation (implanted animals that were subjected to anesthesia but no VNS) three hours before LPS administration. Out of the 7 stimulated mice, VNS produced a decrease in HR in 4 animals, of which 2 exhibited more that 50% decrease in serum TNF compared to sham-stimulated controls and animals with no HR response (Fig 5B). Mice that lacked a physiological response had TNF levels comparable to sham-stimulated controls (Fig 5B).

**Fig 5.**
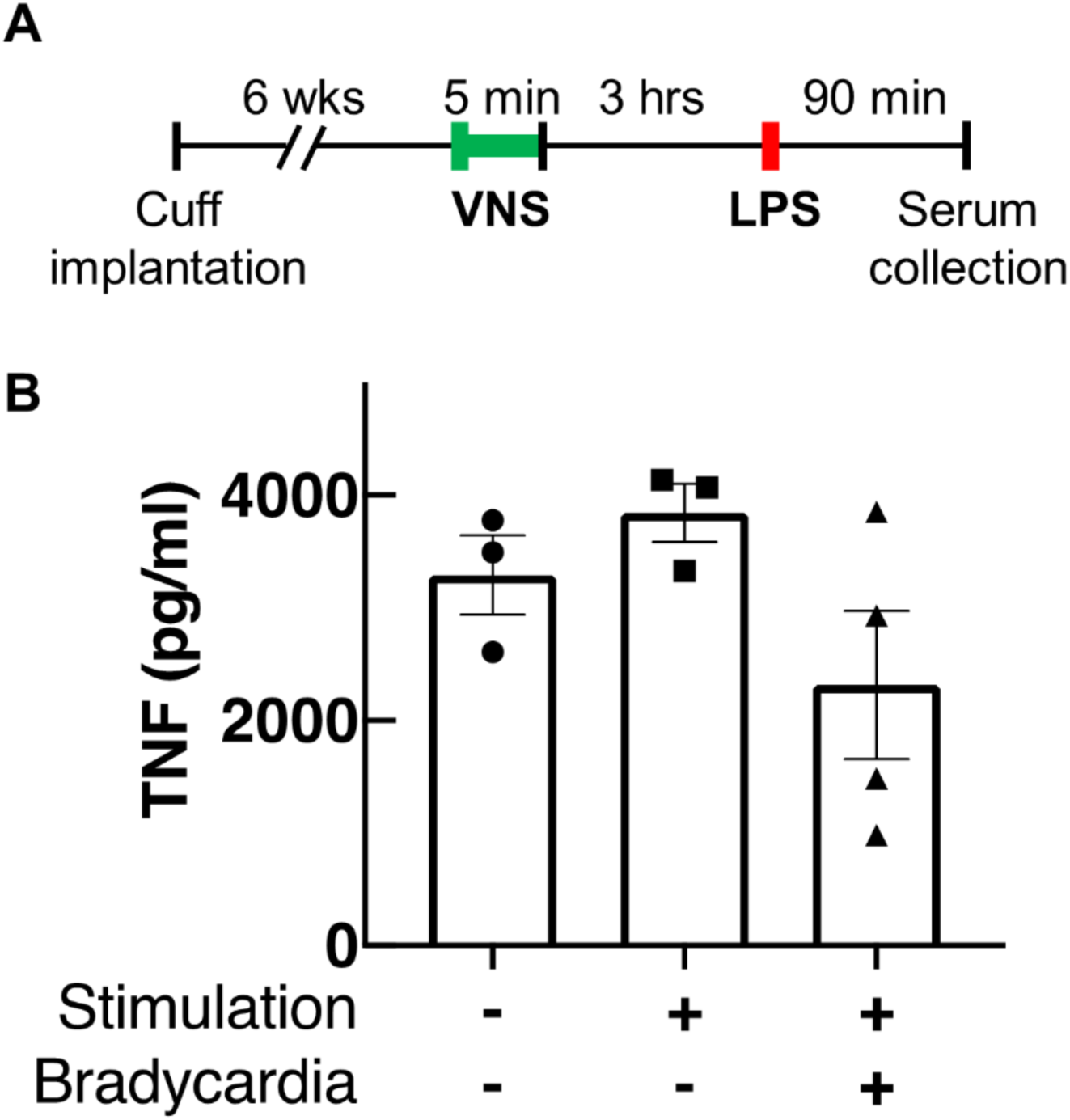
One-time VNS through long-term implants in LPS endotoxemia. (A) Ten mice with 6-week old implants received either electrical or sham VNS for 5 minutes using previously reported parameters (intensity at HRT, PW 250 μs, frequency 10 Hz). LPS was administered 3 hours later, and blood was collected 90 min post LPS-injection. (B) Serum TNF levels from mice that received sham stimulation (left bar), VNS without HR response (middle bar) and VNS eliciting bradycardia (right bar). Data shown as mean ± SEM. *P* = 0.167, one-way ANOVA.

## Discussion

Vagus nerve stimulation (VNS) is an emerging bioelectronic treatment with possible applications in many chronic diseases. However, its translational potential is hindered by the lack of a reliable long-term VNS implant in mice—the preferred species in the preclinical study of human diseases [62]. Development of a simple, well-characterized and reliable long-term VNS interface in mice will allow for standardized assessment of long-term VNS efficacy, as well as potential adverse effects, in various models of chronic disease. It will also allow mechanistic studies of autonomic tone alterations that might accompany long-term VNS. Here, we describe a surgical approach to permanently implant a micro-cuff electrode onto the mouse cervical VN and provide a systematic method to assess its functionality over time as well as standardize stimulation dosing. Our data demonstrate a robust and reproducible interface with the VN capable of producing characteristic physiological responses up to 90 days post-implantation, while causing no large-scale axonal damage.

To maximize the applicability of our tool in various research programs carried out by teams with different areas of expertise, we tried to accomplish 2 goals: ease of assembly and use, and reproducibility. Assembly of the implant makes use of only off-the-shelf supplies and materials, including commercially available micro-cuffs and common physiological sensors. Determining HRT requires the use of a simple rodent heart monitor. The surgical technique, that reflects the aggregate experience of 3 research groups, was refined and simplified over the course of several animal cohorts. Finally, validation of the longitudinal performance of this implant by 2 research groups supports the reproducibility of this method when exercised by different investigators.

Previous research using VNS in mouse models of disease has been limited to acute, single-event, stimulation [44, 49, 63]. These studies, although of translational value, provide limited insight into the possible role of VNS in the treatment of chronic conditions. Moreover, acute stimulation studies are carried out under anesthesia, which confounds the results due to the effects of some anesthetics, such as isoflurane, on decreasing vagal tone [64, 65] and suppressing the immune response [66]. In this study, we established the feasibility of a long-term vagus interface in mice capable of producing stimulus-elicited responses for at least 4 weeks, a period that allows for therapeutic assessment in many models of chronic disease. We also showed that this approach can be used to deliver dose-controlled VNS in conscious animals, hence avoiding repeated exposure to anesthetics during the course of long-term stimulation, while ensuring consistent stimulation longitudinally and across different animals (S1 Movie). VNS in 2 awake behaving animals produced comparable changes in HR as those seen in anesthetized animals and implant longevity in those 2 animals ranged between 2 and 3 weeks.

VNS causes reduction in HR, mainly through activation of efferent cardio-inhibitory fibers [67]. In a functional neural interface, increasing stimulation intensity leads to recruitment of more fibers, and hence a larger stimulus-evoked response [68]. This is consistent with our observation wherein changes in stimulation intensity resulted in a dose-dependent decrease in heart rate (Fig 2B). Additionally, higher stimulation intensities lead to the ordered recruitment of different fiber types according to size: from large A, to intermediate-size B, to small C fibers [69]. The variable changes in breathing rate we observed at different stimulation intensities (Figs 2A and 3A) can be explained by activation of either A or C fibers, which differentially affect breathing in mice [59]. In addition, a functional nerve interface exhibits a characteristic relationship between intensity and pulse width (strength-duration): to produce a response, lower stimulation intensities are required at longer pulse widths, and vice versa [70]. Our long-term implants produced HR responses at lower intensities with longer pulse widths, a relationship that remained consistent with time (Fig 3C). These findings indicate a robust electrode-tissue interface with the VN.

An effective approach to delivering long-term VNS must include standardized methods for verifying implant functionality and controlling stimulation dose over the course of treatment. To evaluate electrode performance across time, we used HRT as a quick and accessible measure of fiber recruitment in real-time [71]. Initial threshold values were variable among animals and increased over the first week post-implantation in almost all mice (Fig 3B). The variation in baseline thresholds at the time of implantation was not explained by differences in pre-implantation impedance and is likely due to variability in electrode placement, which affects fiber engagement. The gradual increase in HRT over time can be attributed to fibrotic encapsulation of electrode surfaces, a process that evolves over days to weeks and is known to reduce the efficacy of implanted neural interfaces by increasing tissue resistivity [60, 72-74]. Encapsulation also increases the distance between the nerve and the electrode contact surface. This explains why longer PWs were more effective in eliciting HRT in aged implants, as the effect of distance on activation threshold is weaker for long pulses than for short pulses [75]. In addition to assessing performance, HRT values were used as a method to estimate individualized stimulation doses. Since animals exhibit variable threshold values at baseline and across time, employing fixed parameters would result in variable fiber recruitment, inconsistent therapeutic dosing and, possibly, undesirable off-target effects. This is of particular importance when VNS is delivered therapeutically, wherein standardized doses are desirable within and in between animals, and across time. Previous reports from rat and large animal models implemented a similar approach to adjust stimulation intensity using respiratory twitching [23, 76], and heart rate [44, 77, 78] to standardize stimulation protocols, but not on a dose-by-dose basis. Notably, electrical impedance, which is commonly used as a measure of electrode integrity and performance [79], did not correlate well with changes in HRT on an individual implant basis as implants aged (Fig 3E). In fact, in cohort 2 animals, impedance values tended to decrease over the first two weeks as HRT increased (Figs 3B and 3D). For these reasons, we chose to rely on HRT values as a reliable indicator of electrode integrity and a method to estimate stimulation dosage [80].

Implant longevity can be influenced by abiotic factors [51, 74], among which are lead breakage and electrode degradation. We observed 3 cases of lead wire breakage out of 9 non-functional implants in cohorts 1-3. In all 3 cases, breakage occurred at the junction between the lead wire and electrode, exposing a mechanically weak point that should be reinforced during manufacturing. Neural electrodes can degrade with long-term use [54]. This process is accelerated at stimulation intensities that exceed the electrode’s ability to transfer charge without undergoing irreversible damage (charge injection capacity, CIC) [54, 56]. The damage imparted by exceeding this threshold is not limited to the electrode but can affect the nerve as well [81]. Although we did not examine explanted electrodes for morphological or electrochemical changes, we did take note of platinum-iridium’s maximum charge capacity [55], which was not exceeded in most animals (Fig 3B, S2 Fig). This becomes important when VNS is administered therapeutically in chronic models, wherein stimulation protocols may be employed on a daily or hourly basis for several weeks. Standardizing stimulation dose could prevent exceeding CIC while still delivering therapeutic stimulation. Mechanical forces generated by chronic cuffing can also damage nerves [82-84]. Our histological analysis revealed generally healthy nerves with no obvious fiber loss or axonal fragmentation. However, we interpret this preliminary analysis with caution due to the lack of small-fiber detail in these relatively thick sections (50 μm). Future studies will include higher resolution images with same side non-cuffed controls. Apart from electrode- and tissue-related factors, we found that surgical proficiency contributed greatly to implant success, which is evident from the increase in successful implantations of cohorts 1-3 (Table 1) completed over several months. Over time, fine adjustments to electrode placement, surgical approach, and post-surgical care lead to higher success rates.

The potential wide therapeutic applicability of VNS lies in part in its ability to modulate inflammation, a process thought to underlie the pathogenesis of many common diseases [27-30]. This effect was first described in a model of LPS endotoxemia, wherein acute VNS suppressed TNF release [6], and has since been established as a functional read-out of the anti-inflammatory action of VNS [42]. For these reasons, we sought to determine whether several weeks-old long-term implants could reproduce that effect. We observed a considerable decrease (> 50%) in serum TNF in 2 out of 4 animals with functional cuffs compared to sham stimulation controls. The lack of an anti-inflammatory response in the other 2 animals could have several explanations. Chronic cuffing of the VN might suppress efferent signaling, including VNS anti-inflammatory actions [84], which could be due to chronic compression injury to the nerve [85, 86]. We did not observe gross axonal damage in the cuffed vagus nerves of those animals (Fig 4B), even though loss of smaller fibers cannot be ruled out as mentioned previously. Alternatively, those 2 animals could simply be non-responders seen often in acute studies [42, 44]. Our data also suggest that the presence of a heart rate response in an implant does not necessarily predict engagement of the anti-inflammatory pathway, which is consistent with previous findings in acute studies [44, 87]. Nevertheless, these findings provide evidence that 6-week-old implants can still control acute inflammation.

We have developed a surgical approach to implant micro-cuff electrodes onto the mouse cervical VN to deliver chronic electrostimulation. Our findings demonstrate a stable and robust interface with the VN capable of producing stimulus-evoked responses for at least 4 weeks. We also provide a method to standardized stimulation doses between animals and across time. Given the wide range of illnesses studied in the mouse, this implant methodology provides a previously unavailable tool for rapid screening of long-term VNS in disease models that could potentially introduce novel therapies to many common and rare diseases.

## Acknowledgments

The authors would like to thank Umair Ahmed, Jacquelyn Tomaio, Khaled Qanud, Arielle Gabalski, Shubham Debnath, Todd Levy, Maria Lopez, Joanne Peragine, Loren Reith, and Moon Rob for their assistance in experiments, electrodes, and data analysis. We also thank Tom Coleman and Kevin Tracey for their support and thoughtful discussions. This work was supported in part by DARPA (N66001-17-2-4010), the BRAIN Initiative (NS107616), NICHD (HD088411), NIDCD (DC12557), a Howard Hughes Medical Institute Faculty Scholarship (R.C.F.) and DARPA Biological Technologies Office Targeted Neuroplasticity Training (TNT) HR0011-17-2-0051 (C.W.).

## Author contributions

Conceived and developed model, IM, JH, EP, RF, CW, SZ, YA; designed and performed experiments, IM, NJ, AA, JH; analyzed and interpreted data, IM, YC, SZ; prepared figures, IM, SZ, JH; drafted manuscript, IM; all authors revised and edited manuscript.

---

**S1 Movie.**
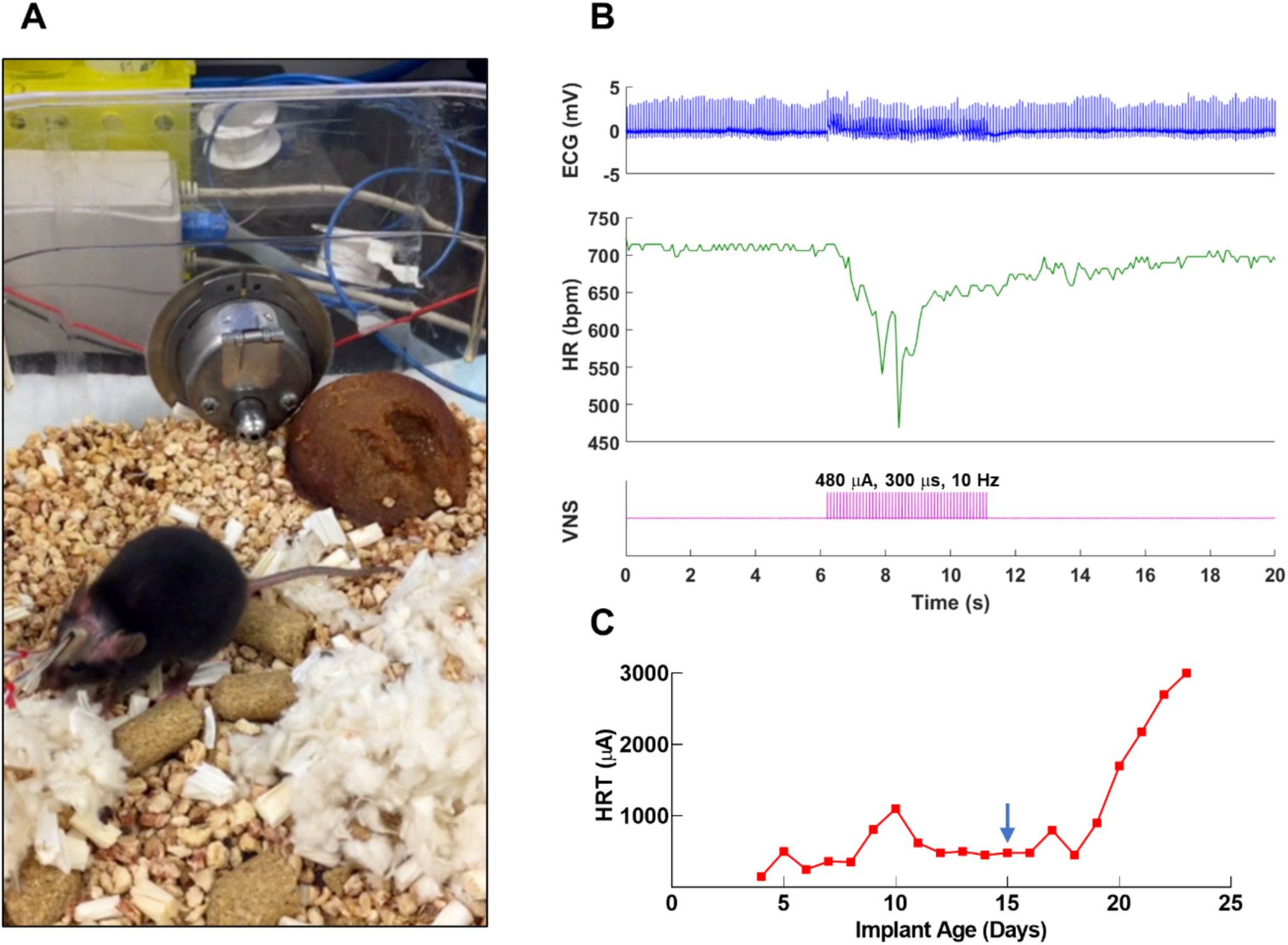
VNS in an awake behaving mouse. (A) Video clip showing an awake behaving mouse with a long-term VN implant and ECG leads, connected to a commutator and receiving VNS on post-implant day 15. The screen shows HR (green trace) and a stimulation event (purple trace). VNS occurs at the 23 s time point. (B) Physiological traces from the stimulation event captured on the video clip. (C) HRT values vs. implant age for mouse shown in video clip. Arrow indicates day on which testing shown in panel B occurred. **Link to video:** https://1drv.ms/u/s!AhZzF8JVQClYgvE_zuKSATPpabHM_Q?e=gbM5AM

**S2 Fig.**
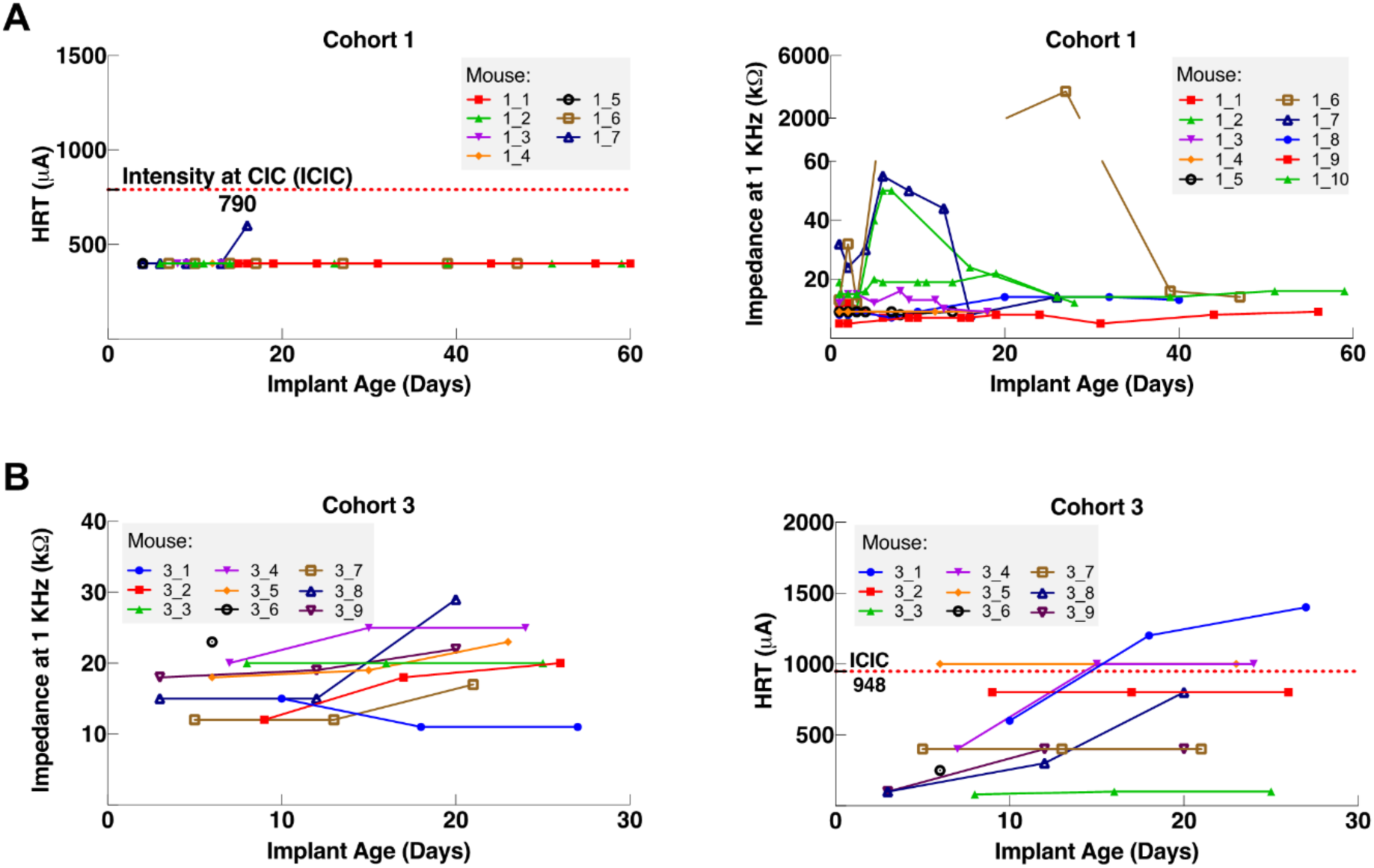
Longitudinal changes in heart rate threshold and electrical impedance in long-term implants in mice from cohorts 1 and 3. (A) HRT values vs. implant age (left panel), and impedance at 1KHz (right panel) in 8 mice from cohort 1 and (B) 9 mice from cohort 3. The horizontal red dotted line indicates intensity at maximum charge injection capacity (ICIC) calculated for the micro-cuff electrodes.

**S3 Fig.**
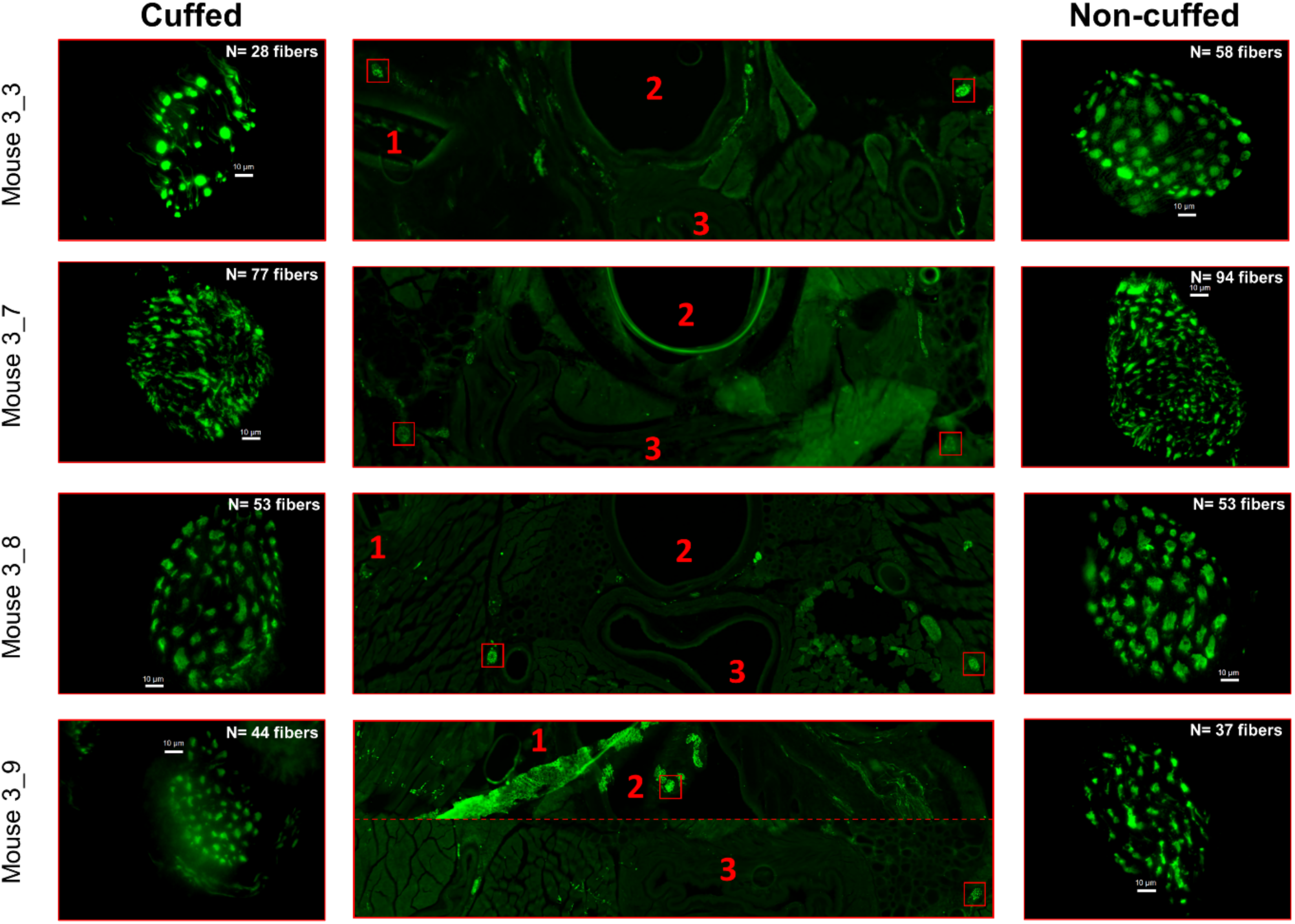
Large nerve fibers in cuffed and non-cuffed VN. Cross-sections of the neck immediately rostral to the cuff (middle panels) from 4 mice with a 6-week old implant stained for axons (neurofilament, green) and showing the following landmarks: (1) rostral cuff margin; (2) trachea; (3) esophagus. The cuffed (left) and non-cuffed (right) VNs were imaged under 100X magnification and fibers larger than 1.5 μm were counted. Scale bars are 10 μm.

## References

1. Berthoud HR, Neuhuber WL. Functional and chemical anatomy of the afferent vagal system. Auton Neurosci. 2000;85(1-3):1–17.

2. Pavlov VA, Chavan SS, Tracey KJ. Molecular and Functional Neuroscience in Immunity. Annu Rev Immunol. 2018;36:783–812.

3. Pavlov VA, Chavan SS, Tracey KJ. Bioelectronic Medicine: From Preclinical Studies on the Inflammatory Reflex to New Approaches in Disease Diagnosis and Treatment. Cold Spring Harb Perspect Med. 2020;10(3).

4. Tracey KJ. The inflammatory reflex. Nature. 2002;420(6917):853–9.

5. Tracey KJ. Reflex control of immunity. Nat Rev Immunol. 2009;9(6):418–28.

6. Borovikova LV, Ivanova S, Zhang M, Yang H, Botchkina GI, Watkins LR, et al. Vagus nerve stimulation attenuates the systemic inflammatory response to endotoxin. Nature. 2000;405(6785):458–62.

7. Rosas-Ballina M, Ochani M, Parrish WR, Ochani K, Harris YT, Huston JM, et al. Splenic nerve is required for cholinergic antiinflammatory pathway control of TNF in endotoxemia. Proceedings of the National Academy of Sciences. 2008;105(31):11008–13.

8. Rosas-Ballina M, Olofsson PS, Ochani M, Valdés-Ferrer SI, Levine YA, Reardon C, et al. Acetylcholine-synthesizing T cells relay neural signals in a vagus nerve circuit. Science. 2011;334(6052):98–101.

9. A randomized controlled trial of chronic vagus nerve stimulation for treatment of medically intractable seizures. The Vagus Nerve Stimulation Study Group. Neurology. 1995;45(2):224–30.

10. Rush AJ, George MS, Sackeim HA, Marangell LB, Husain MM, Giller C, et al. Vagus nerve stimulation (VNS) for treatment-resistant depressions: a multicenter study. Biol Psychiatry. 2000;47(4):276–86.

11. Tyler R, Cacace A, Stocking C, Tarver B, Engineer N, Martin J, et al. Vagus Nerve Stimulation Paired with Tones for the Treatment of Tinnitus: A Prospective Randomized Double-blind Controlled Pilot Study in Humans. Sci Rep. 2017;7(1):11960.

12. Vanneste S, Martin J, Rennaker RL, 2nd, Kilgard MP. Pairing sound with vagus nerve stimulation modulates cortical synchrony and phase coherence in tinnitus: An exploratory retrospective study. Sci Rep. 2017;7(1):17345.

13. Engineer ND, Kimberley TJ, Prudente CN, Dawson J, Tarver WB, Hays SA. Targeted Vagus Nerve Stimulation for Rehabilitation After Stroke. Front Neurosci. 2019;13:280.

14. Kimberley TJ, Pierce D, Prudente CN, Francisco GE, Yozbatiran N, Smith P, et al. Vagus Nerve Stimulation Paired With Upper Limb Rehabilitation After Chronic Stroke. Stroke. 2018;49(11):2789–92.

15. Merrill CA, Jonsson MA, Minthon L, Ejnell H, H CsS, Blennow K, et al. Vagus nerve stimulation in patients with Alzheimer’s disease: Additional follow-up results of a pilot study through 1 year. J Clin Psychiatry. 2006;67(8):1171–8.

16. Sjogren MJ, Hellstrom PT, Jonsson MA, Runnerstam M, Silander HC, Ben-Menachem E. Cognition-enhancing effect of vagus nerve stimulation in patients with Alzheimer’s disease: a pilot study. J Clin Psychiatry. 2002;63(11):972–80.

17. De Ferrari GM, Stolen C, Tuinenburg AE, Wright DJ, Brugada J, Butter C, et al. Long-term vagal stimulation for heart failure: Eighteen month results from the NEural Cardiac TherApy foR Heart Failure (NECTAR-HF) trial. Int J Cardiol. 2017;244:229–34.

18. Huang WA, Shivkumar K, Vaseghi M. Device-based autonomic modulation in arrhythmia patients: the role of vagal nerve stimulation. Curr Treat Options Cardiovasc Med. 2015;17(5):379.

19. Nasi-Er BG, Wenhui Z, HuaXin S, Xianhui Z, Yaodong L, Yanmei L, et al. Vagus nerve stimulation reduces ventricular arrhythmias and increases ventricular electrical stability. Pacing Clin Electrophysiol. 2019;42(2):247–56.

20. Stavrakis S, Humphrey MB, Scherlag BJ, Hu Y, Jackman WM, Nakagawa H, et al. Low-level transcutaneous electrical vagus nerve stimulation suppresses atrial fibrillation. J Am Coll Cardiol. 2015;65(9):867–75.

21. Yamaguchi N, Yamakawa K, Rajendran PS, Takamiya T, Vaseghi M. Antiarrhythmic effects of vagal nerve stimulation after cardiac sympathetic denervation in the setting of chronic myocardial infarction. Heart Rhythm. 2018;15(8):1214–22.

22. Ntiloudi D, Qanud K, Tomaio JN, Giannakoulas G, Al-Abed Y, Zanos S. Pulmonary arterial hypertension: the case for a bioelectronic treatment. Bioelectron Med. 2019;5:20.

23. Yoshida K, Saku K, Kamada K, Abe K, Tanaka-Ishikawa M, Tohyama T, et al. Electrical Vagal Nerve Stimulation Ameliorates Pulmonary Vascular Remodeling and Improves Survival in Rats With Severe Pulmonary Arterial Hypertension. JACC Basic Transl Sci. 2018;3(5):657–71.

24. Koopman FA, Chavan SS, Miljko S, Grazio S, Sokolovic S, Schuurman PR, et al. Vagus nerve stimulation inhibits cytokine production and attenuates disease severity in rheumatoid arthritis. Proc Natl Acad Sci U S A. 2016;113(29):8284–9.

25. Bonaz B, Sinniger V, Hoffmann D, Clarencon D, Mathieu N, Dantzer C, et al. Chronic vagus nerve stimulation in Crohn’s disease: a 6-month follow-up pilot study. Neurogastroenterol Motil. 2016;28(6):948–53.

26. Aranow C. Engaging the Cholinergic Anti-Inflammatory Pathway By Stimulating the Vagus Nerve Reduces Pain and Fatigue in Patients with SLE. 2018 ACR/ARHP Annual Meeting; 2018: ACR.

27. Couzin-Frankel J. Inflammation bares a dark side. Science. 2010;330(6011):1621.

28. Furman D, Campisi J, Verdin E, Carrera-Bastos P, Targ S, Franceschi C, et al. Chronic inflammation in the etiology of disease across the life span. Nat Med. 2019;25(12):1822–32.

29. Slavich GM. Understanding inflammation, its regulation, and relevance for health: a top scientific and public priority. Brain Behav Immun. 2015;45:13–4.

30. Strowig T, Henao-Mejia J, Elinav E, Flavell R. Inflammasomes in health and disease. Nature. 2012;481(7381):278–86.

31. Annoni EM, Van Helden D, Guo Y, Levac B, Libbus I, KenKnight BH, et al. Chronic Low-Level Vagus Nerve Stimulation Improves Long-Term Survival in Salt-Sensitive Hypertensive Rats. Front Physiol. 2019;10:25.

32. Beaumont E, Wright GL, Southerland EM, Li Y, Chui R, KenKnight BH, et al. Vagus nerve stimulation mitigates intrinsic cardiac neuronal remodeling and cardiac hypertrophy induced by chronic pressure overload in guinea pig. Am J Physiol Heart Circ Physiol. 2016;310(10):H1349–59.

33. Chinda K, Tsai WC, Chan YH, Lin AY, Patel J, Zhao Y, et al. Intermittent left cervical vagal nerve stimulation damages the stellate ganglia and reduces the ventricular rate during sustained atrial fibrillation in ambulatory dogs. Heart Rhythm. 2016;13(3):771–80.

34. Farrand AQ, Helke KL, Gregory RA, Gooz M, Hinson VK, Boger HA. Vagus nerve stimulation improves locomotion and neuronal populations in a model of Parkinson’s disease. Brain Stimul. 2017;10(6):1045–54.

35. Li M, Zheng C, Sato T, Kawada T, Sugimachi M, Sunagawa K. Vagal nerve stimulation markedly improves long-term survival after chronic heart failure in rats. Circulation. 2004;109(1):120–4.

36. Meyers EC, Solorzano BR, James J, Ganzer PD, Lai ES, Rennaker RL, 2nd, et al. Vagus Nerve Stimulation Enhances Stable Plasticity and Generalization of Stroke Recovery. Stroke. 2018;49(3):710–7.

37. Nuntaphum W, Pongkan W, Wongjaikam S, Thummasorn S, Tanajak P, Khamseekaew J, et al. Vagus nerve stimulation exerts cardioprotection against myocardial ischemia/reperfusion injury predominantly through its efferent vagal fibers. Basic Res Cardiol. 2018;113(4):22.

38. Yamakawa K, Rajendran PS, Takamiya T, Yagishita D, So EL, Mahajan A, et al. Vagal nerve stimulation activates vagal afferent fibers that reduce cardiac efferent parasympathetic effects. Am J Physiol Heart Circ Physiol. 2015;309(9):H1579–90.

39. Bryda EC. The Mighty Mouse: the impact of rodents on advances in biomedical research. Mo Med. 2013;110(3):207–11.

40. Perlman RL. Mouse models of human disease: An evolutionary perspective. Evol Med Public Health. 2016;2016(1):170–6.

41. Bansal V, Ryu SY, Lopez N, Allexan S, Krzyzaniak M, Eliceiri B, et al. Vagal stimulation modulates inflammation through a ghrelin mediated mechanism in traumatic brain injury. Inflammation. 2012;35(1):214–20.

42. Caravaca AS, Gallina AL, Tarnawski L, Tracey KJ, Pavlov VA, Levine YA, et al. An Effective Method for Acute Vagus Nerve Stimulation in Experimental Inflammation. Front Neurosci. 2019;13:877.

43. de Lucas-Cerrillo AM, Maldifassi MC, Arnalich F, Renart J, Atienza G, Serantes R, et al. Function of partially duplicated human alpha77 nicotinic receptor subunit CHRFAM7A gene: potential implications for the cholinergic anti-inflammatory response. J Biol Chem. 2011;286(1):594–606.

44. Huffman WJ, Subramaniyan S, Rodriguiz RM, Wetsel WC, Grill WM, Terrando N. Modulation of neuroinflammation and memory dysfunction using percutaneous vagus nerve stimulation in mice. Brain Stimul. 2019;12(1):19–29.

45. Huston JM, Ochani M, Rosas-Ballina M, Liao H, Ochani K, Pavlov VA, et al. Splenectomy inactivates the cholinergic antiinflammatory pathway during lethal endotoxemia and polymicrobial sepsis. J Exp Med. 2006;203(7):1623–8.

46. Meneses G, Bautista M, Florentino A, Diaz G, Acero G, Besedovsky H, et al. Electric stimulation of the vagus nerve reduced mouse neuroinflammation induced by lipopolysaccharide. J Inflamm (Lond). 2016;13:33.

47. Saeed RW, Varma S, Peng-Nemeroff T, Sherry B, Balakhaneh D, Huston J, et al. Cholinergic stimulation blocks endothelial cell activation and leukocyte recruitment during inflammation. J Exp Med. 2005;201(7):1113–23.

48. Shukla S, Fritz JR, Kyaw CA, Valentino SP, Imossi CW, Sarangi SN, et al. Cholinergic Stimulation Improves Hemostasis in a Hemophilia Mouse Model. Blood. 2015;126(23):3528-.

49. The FO, Boeckxstaens GE, Snoek SA, Cash JL, Bennink R, Larosa GJ, et al. Activation of the cholinergic anti-inflammatory pathway ameliorates postoperative ileus in mice. Gastroenterology. 2007;133(4):1219–28.

50. Cuoco FA, Jr., Durand DM. Measurement of external pressures generated by nerve cuff electrodes. IEEE Trans Rehabil Eng. 2000;8(1):35–41.

51. Prasad A, Xue QS, Dieme R, Sankar V, Mayrand RC, Nishida T, et al. Abiotic-biotic characterization of Pt/Ir microelectrode arrays in chronic implants. Front Neuroeng. 2014;7:2.

52. Rydevik B, Lundborg G, Bagge U. Effects of graded compression on intraneural blood blow. An in vivo study on rabbit tibial nerve. J Hand Surg Am. 1981;6(1):3–12.

53. McCreery DB, Agnew WF, Yuen TG, Bullara LA. Damage in peripheral nerve from continuous electrical stimulation: comparison of two stimulus waveforms. Med Biol Eng Comput. 1992;30(1):109–14.

54. Negi S, Bhandari R, Rieth L, Van Wagenen R, Solzbacher F. Neural electrode degradation from continuous electrical stimulation: comparison of sputtered and activated iridium oxide. J Neurosci Methods. 2010;186(1):8–17.

55. Cogan SF. Neural stimulation and recording electrodes. Annu Rev Biomed Eng. 2008;10:275–309.

56. Merrill DR, Bikson M, Jefferys JG. Electrical stimulation of excitable tissue: design of efficacious and safe protocols. J Neurosci Methods. 2005;141(2):171–98.

57. Crosby K, Simendinger J, Grange C, Ferrante M, Bernier T, Stanen C. Immunohistochemistry protocol for paraffin-embedded tissue section-advertisement. Cell Signal Technol. 2016.

58. Agostoni E, Chinnock JE, De Daly MB, Murray JG. Functional and histological studies of the vagus nerve and its branches to the heart, lungs and abdominal viscera in the cat. J Physiol. 1957;135(1):182–205.

59. Chang RB, Strochlic DE, Williams EK, Umans BD, Liberles SD. Vagal Sensory Neuron Subtypes that Differentially Control Breathing. Cell. 2015;161(3):622–33.

60. Anderson JM, Rodriguez A, Chang DT. Foreign body reaction to biomaterials. Semin Immunol. 2008;20(2):86–100.

61. Tyler DJ, Durand DM. Chronic response of the rat sciatic nerve to the flat interface nerve electrode. Ann Biomed Eng. 2003;31(6):633–42.

62. Vandenbergh JG. The House Mouse in Biomedical Research. Sourcebook of Models for Biomedical Research: Springer; 2008. p. 187–90.

63. Ji H, Rabbi MF, Labis B, Pavlov VA, Tracey KJ, Ghia JE. Central cholinergic activation of a vagus nerve-to-spleen circuit alleviates experimental colitis. Mucosal Immunol. 2014;7(2):335–47.

64. Marano G, Grigioni M, Tiburzi F, Vergari A, Zanghi F. Effects of isoflurane on cardiovascular system and sympathovagal balance in New Zealand white rabbits. J Cardiovasc Pharmacol. 1996;28(4):513–8.

65. Picker O, Scheeren TW, Arndt JO. Inhalation anaesthetics increase heart rate by decreasing cardiac vagal activity in dogs. Br J Anaesth. 2001;87(5):748–54.

66. Cruz FF, Rocco PR, Pelosi P. Anti-inflammatory properties of anesthetic agents. Crit Care. 2017;21(1):67.

67. Buschman HP, Storm CJ, Duncker DJ, Verdouw PD, van der Aa HE, van der Kemp P. Heart rate control via vagus nerve stimulation. Neuromodulation. 2006;9(3):214–20.

68. Zaaimi B, Grebe R, Wallois F. Animal model of the short-term cardiorespiratory effects of intermittent vagus nerve stimulation. Auton Neurosci. 2008;143(1-2):20–6.

69. Blair E, Erlanger J. A comparison of the characteristics of axons through their individual electrical responses. American Journal of Physiology-Legacy Content. 1933;106(3):524–64.

70. Geddes LA, Bourland JD. The strength-duration curve. IEEE Trans Biomed Eng. 1985;32(6):458–9.

71. Yoo PB, Liu H, Hincapie JG, Ruble SB, Hamann JJ, Grill WM. Modulation of heart rate by temporally patterned vagus nerve stimulation in the anesthetized dog. Physiol Rep. 2016;4(2).

72. Grill WM, Mortimer JT. Electrical properties of implant encapsulation tissue. Ann Biomed Eng. 1994;22(1):23–33.

73. Farah S, Doloff JC, Muller P, Sadraei A, Han HJ, Olafson K, et al. Long-term implant fibrosis prevention in rodents and non-human primates using crystallized drug formulations. Nat Mater. 2019;18(8):892–904.

74. Vasudevan S, Patel K, Welle C. Rodent model for assessing the long term safety and performance of peripheral nerve recording electrodes. J Neural Eng. 2017;14(1):016008.

75. Grill WM, Mortimer JT. Stimulus waveforms for selective neural stimulation. IEEE Engineering in Medicine and Biology Magazine. 1995;14(4):375–85.

76. Nishizaki A, Sakamoto K, Saku K, Hosokawa K, Sakamoto T, Oga Y, et al. Optimal Titration Is Important to Maximize the Beneficial Effects of Vagal Nerve Stimulation in Chronic Heart Failure. J Card Fail. 2016;22(8):631–8.

77. Chapleau MW, Rotella DL, Reho JJ, Rahmouni K, Stauss HM. Chronic vagal nerve stimulation prevents high-salt diet-induced endothelial dysfunction and aortic stiffening in stroke-prone spontaneously hypertensive rats. Am J Physiol Heart Circ Physiol. 2016;311(1):H276–85.

78. Yu H, Tang M, Yu J, Zhou X, Zeng L, Zhang S. Chronic vagus nerve stimulation improves left ventricular function in a canine model of chronic mitral regurgitation. J Transl Med. 2014;12:302.

79. Straka MM, Shafer B, Vasudevan S, Welle C, Rieth L. Characterizing Longitudinal Changes in the Impedance Spectra of In-Vivo Peripheral Nerve Electrodes. Micromachines (Basel). 2018;9(11).

80. Qing KY, Wasilczuk KM, Ward MP, Phillips EH, Vlachos PP, Goergen CJ, et al. B fibers are the best predictors of cardiac activity during Vagus nerve stimulation: Qing, vagal B fiber activation and cardiac effects. Bioelectron Med. 2018;4:5.

81. Cogan SF, Ludwig KA, Welle CG, Takmakov P. Tissue damage thresholds during therapeutic electrical stimulation. J Neural Eng. 2016;13(2):021001.

82. Agnew WF, McCreery DB. Considerations for safety with chronically implanted nerve electrodes. Epilepsia. 1990;31 Suppl 2:S27–32.

83. Larsen JO, Thomsen M, Haugland M, Sinkjaer T. Degeneration and regeneration in rabbit peripheral nerve with long-term nerve cuff electrode implant: a stereological study of myelinated and unmyelinated axons. Acta Neuropathol. 1998;96(4):365–78.

84. Somann JP, Albors GO, Neihouser KV, Lu KH, Liu Z, Ward MP, et al. Chronic cuffing of cervical vagus nerve inhibits efferent fiber integrity in rat model. J Neural Eng. 2018;15(3):036018.

85. Nitz AJ, Dobner JJ, Matulionis DH. Pneumatic tourniquet application and nerve integrity: motor function and electrophysiology. Exp Neurol. 1986;94(2):264–79.

86. Szabo RM, Sharkey NA. Response of peripheral nerve to cyclic compression in a laboratory rat model. J Orthop Res. 1993;11(6):828–33.

87. Huston JM, Gallowitsch-Puerta M, Ochani M, Ochani K, Yuan R, Rosas-Ballina M, et al. Transcutaneous vagus nerve stimulation reduces serum high mobility group box 1 levels and improves survival in murine sepsis. Crit Care Med. 2007;35(12):2762–8.

